# Improved Allele Frequencies in gnomAD through Local Ancestry Inference

**DOI:** 10.1101/2024.10.30.620961

**Authors:** Pragati Kore, Michael W. Wilson, Grace Tiao, Katherine Chao, Philip W. Darnowsky, Nick Watts, Jessica Honorato Mauer, Samantha M. Baxter, Genome Aggregation Database Consortium, Heidi L Rehm, Mark J. Daly, Konrad J. Karczewski, Elizabeth G. Atkinson

## Abstract

The Genome Aggregation Database (gnomAD) is a foundational resource for allele frequency data, widely used in genomic research and clinical interpretation. However, traditional estimates rely on individual-level genetic ancestry groupings that may obscure variation in recently admixed populations. To improve resolution, we applied local ancestry inference (LAI) to over 27 million variants in two admixed groups: Admixed American (n = 7,612) and African/African American (n = 20,250), deriving ancestry-specific allele frequencies. We show that 78.5% and 85.1% of variants in these groups, respectively, exhibit at least a twofold difference in ancestry-specific frequencies. Moreover, 81.49% of variants with LAI information would be assigned a higher gnomAD-wide maximum frequency after incorporating LAI, potentially altering clinical interpretations. This LAI-informed release reveals clinically relevant frequency differences that are masked in aggregate estimates and may support reclassifying some variants from Uncertain Significance to Benign or Likely Benign.

## INTRODUCTION

The Genome Aggregation Database (gnomAD) is one of the most widely used resources in the genomics community^1^, offering comprehensive data on allele frequencies across diverse genetic ancestry groups. With its vast catalog of genetic variation derived from hundreds of thousands of individuals, gnomAD has become an invaluable tool for researchers and clinicians, aiding in the interpretation of genetic variants and their potential implications for human health^2,3^. By providing allele frequency data across diverse genetic ancestry groups, gnomAD helps in the interpretation of the impact of rare and common genetic variants, making it a cornerstone of modern genetic research.

Allele frequencies are a critical component of genomics research and clinical genetics, serving as a key metric for understanding the distribution of genetic variants within and between genetic ancestry groups^3^. These frequencies are used to assess the pathogenicity of variants, guide the design of genetic data generation platforms, and inform the development of therapeutic strategies. In particular, allele frequencies stratified by genetic ancestry groups are essential for identifying genetic risks that are enriched in particular groups and understanding the genetic architecture of complex traits. As such, gnomAD’s genetic ancestry group allele frequencies have been pivotal in advancing our understanding of human genetic variation and its impact on health and disease.

Despite the utility of gnomAD’s allele frequencies, there are limitations inherent in the way these frequencies are computed and how genetic ancestry is represented. While genetic ancestry is truly continuous^4^, the genetic ancestry groups used in gnomAD’s aggregations are, by nature, discrete. Specifically, these groups are defined by assigning individuals a population label defined by global genetic ancestry as trained on reference-panel informed principal components^1^, which can obscure finer-scale genetic differences observed across individuals within a genetic ancestry group^5–8^. This limitation is particularly important when considering groups that have high amounts of genetic diversity or those with recent admixture events, including the gnomAD groups currently identified as ‘Admixed Americans’ and ‘African/African Americans’. Additionally, the use of individual-level groupings does not capture the full spectrum of an individual’s genetic background, which can span multiple continental ancestries.

Local ancestry inference (LAI) offers a solution to some of these limitations by deconvolving allele frequencies based on the specific ancestry of chromosomal segments, rather than entire individuals^9^. By applying LAI, we resolve finer-scale allele frequencies, allowing for better-informed clinical interpretations. This added resolution also increases confidence in classifying variants as benign/likely benign when they are common in any ancestry, thereby lowering the overall burden of variants of uncertain significance across all genetic ancestry groups.

We wish to be clear that in all analyses we exclusively examine genetic ancestry. Assignment of an individual’s genetic ancestry is not equivalent to and does not negate how an individual self-identifies. To refer to the samples included in these analyses, we adhere to the current nomenclature in gnomAD, namely “Admixed American” and “African/African American”. We acknowledge that this terminology is not ideal, however for consistency with broader efforts utilizing these data, and to be clear on the individuals included in these analyses, we retain it here. We look forward to continuing to further refine gnomAD allele frequency presentations in future releases to better reflect the continuous nature of genetic ancestry^5,6^. Throughout this article, we use the term “Amerindigenous” to refer to local ancestry tracts inferred to originate from Indigenous American populations, based on reference haplotypes harmonized from AMR-designated samples in the Human Genome Diversity Project (HGDP) and 1000 Genomes Project Phase 3 (1kGP3). To distinguish between individual-level genetic ancestry groupings (e.g., Admixed American) and ancestry-specific segments within individuals, we refer to frequencies derived from painted tracts as LAI-AMR (Amerindigenous), LAI-AFR (African), and LAI-EUR (European), respectively. Here, we enhance the granularity of gnomAD’s allele frequency data in the inferred Admixed American and African/African American genetic ancestry groups, providing a more accurate representation of genetic variation and its implications for human health.

## RESULTS

### Release of local ancestry-informed frequencies in gnomAD

Here we release local ancestry-informed frequency data for 14,804,206 bi-allelic single nucleotide polymorphisms (SNPs) of at least MAF 0.1% within the Admixed American (n=7,612) and 24,204,574 bi-allelic SNPs within the African/African American genetic ancestry groups (n=20,250), both from gnomAD v3.1. Of the total SNPs in each genetic ancestry group, 9.82% of SNPs in the African/African American group and 11.57% of SNPs in the Admixed American group passed QC and were assigned local ancestry calls. This initial release of gnomAD LAI data contains variants’ estimated alternate allele counts, allele numbers, and allele frequencies partitioned by haplotype-level continental ancestries. These new local ancestry-informed frequencies are released in a downloadable VCF format file and can be explored interactively on the gnomAD browser (https://gnomad.broadinstitute.org/). To access the local ancestry data for a specific variant within the browser, select the “Local Ancestry” tab within the genetic ancestry group frequency table on the variant’s page. If local ancestry data is available for the variant, the ancestral allele count (AC), allele number (AN), and allele frequency (AF) estimates will be displayed (**Supplementary Fig. 1**).

### Ancestry composition of gnomAD African/African Americans and Admixed Americans

Plotting the first two PCs of the full gnomAD collection, we can see the inferred gnomAD Admixed Americans span a large portion of global PC space, with some samples clustering closer to European reference samples, others clustering closer to the African reference samples, and some clustering closer to the East Asian reference samples, indexing the presence of Amerindigenous ancestry^10^ (**Fig. 1a, Supplementary Fig. 2a**). Resolving global ancestry with ADMIXTURE, we again observe that samples classified as Admixed American by gnomAD exhibit heterogeneous genetic ancestry, with individuals containing differing proportions of two or three-way admixture between Amerindigenous (AMR), African (AFR), and European (EUR) haplotypes. After applying a 5% ancestry inclusion threshold, 5% of individuals derive their genetic ancestry primarily from a single continent (homogeneous AMR or EUR or AFR), 60% from two continents (AFR/AMR or AFR/EUR or AMR/EUR), and 35% from three continents (AMR/EUR/AFR) (**Fig. 1b**). Given this composition, we modeled the Admixed American samples with a three-way reference panel (see *Methods*). The African/African American samples primarily lie on a cline between the EUR and AFR poles in PC space, indicating admixture between largely these two genetic ancestry components. Further exploring African/African American samples’ ancestry with ADMIXTURE, we again observed individuals in this group contain primarily AFR and EUR components, with 32% homogeneous AFR individuals present and 68% of samples classified as two-way admixed (**Fig. 1c**). Therefore, we curated a reference panel comprising two-way continental AFR and EUR genetic ancestries for this group.

**Figure 1.**
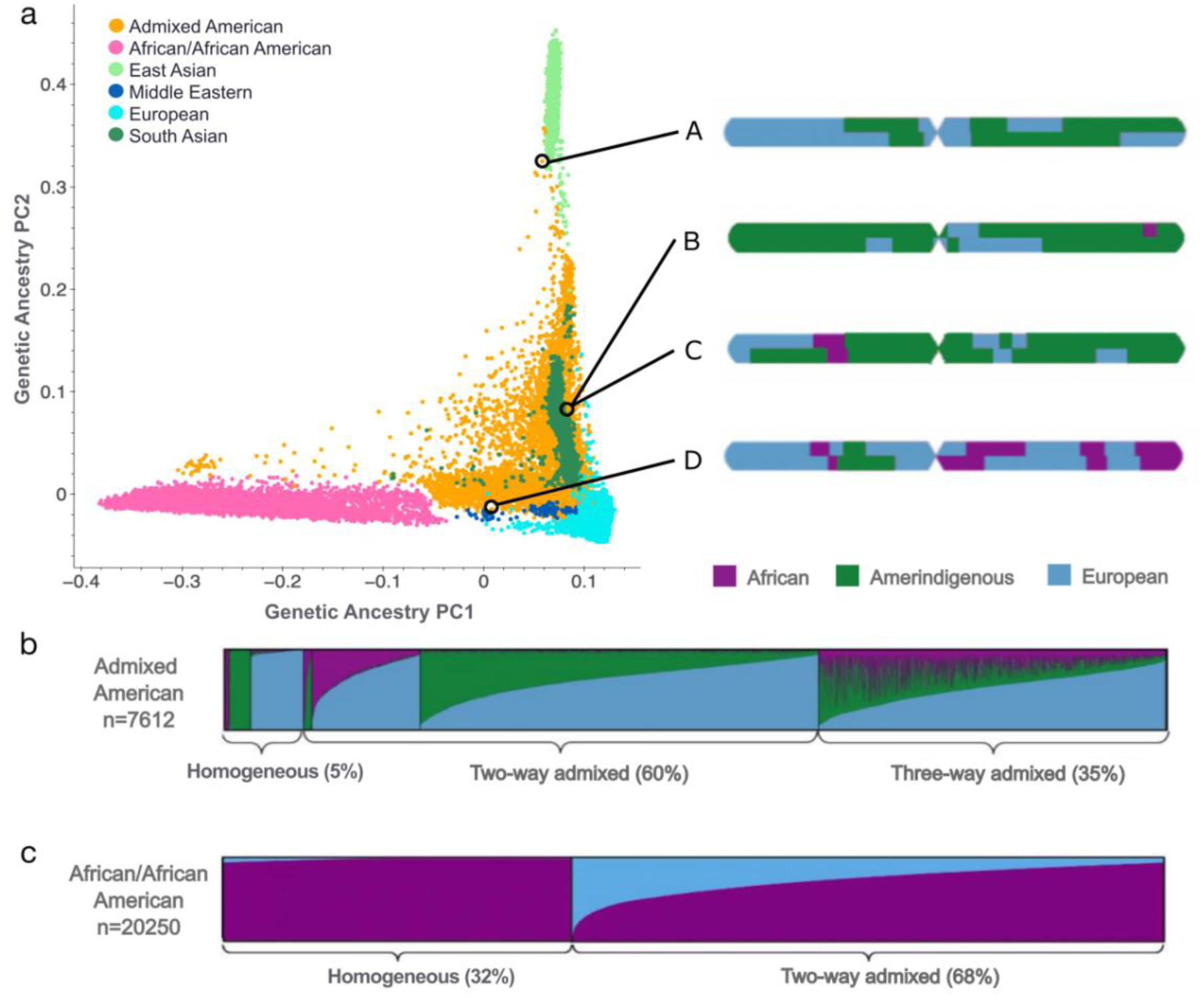
Population structure of the gnomAD Admixed American and African/African American genetic ancestry groups. **a.** PCA plot for all gnomAD samples with the painted karyograms of a single chromosome from an individual within each circle. Note that individuals B and C have strikingly different local ancestry patterns despite appearing in the same location in PC space. **b.** ADMIXTURE plot for the gnomAD Admixed American genetic ancestry group displaying global ancestry proportions. 5% of individuals derive ancestry from a single continent, 60% from two, and 35% from three. **c.** ADMIXTURE plot for the African/African American genetic ancestry group showed admixture primarily from two continents, with 32% homogeneous AFR individuals.

Local ancestry deconvolution provides additional resolution beyond global inference; we observe that even individuals occupying the same point in PC space have markedly different local ancestry patterns (**Fig. 1a**). Notably, even across higher principal components (e.g., PC1 to PC12), individuals ‘b’ and ‘c’ remain closely clustered, indicating that despite differences in their local ancestry, their overall genetic profiles are still similar when considering finer-scale genetic variation (**Supplementary Fig. 2b-g**). With only genetic ancestry group level rather than ancestry-specific allele frequencies, estimates represent a weighted average of haplotype-derived frequencies and can thus be distorted from the more precise ancestry-specific frequencies. This additional resolution can be meaningful for variants of functional importance.

### Functional variant case studies

For example, the gnomAD variant 17-7043011-C-T in *SLC16A11*, associated with type 2 diabetes risk, is elevated within the Admixed American genetic ancestry group^11^ at a 24% allele frequency compared to the gnomAD-wide gnomAD allele frequency of 2% (**Fig. 2c**). Breaking down this variant’s frequency into continental allele frequency estimates, we see in fact that the variant occurs at a 45% allele frequency in LAI-AMR regions and closer to 0% in the LAI-AFR and LAI-EUR regions. Similarly, the gnomAD variant 22-36265860-A-G in *APOL1*, linked with increased susceptibility to focal segmental glomerulosclerosis (FSGS), HIV-associated nephropathy (HIVAN), and hypertensive end-stage kidney disease (ESKD)^12^, shows a 27% allele frequency in LAI-AFR regions of the African/African American genetic ancestry group^13^, compared to a 1% gnomAD-wide frequency (**Fig. 2d**).

**Figure 2:**
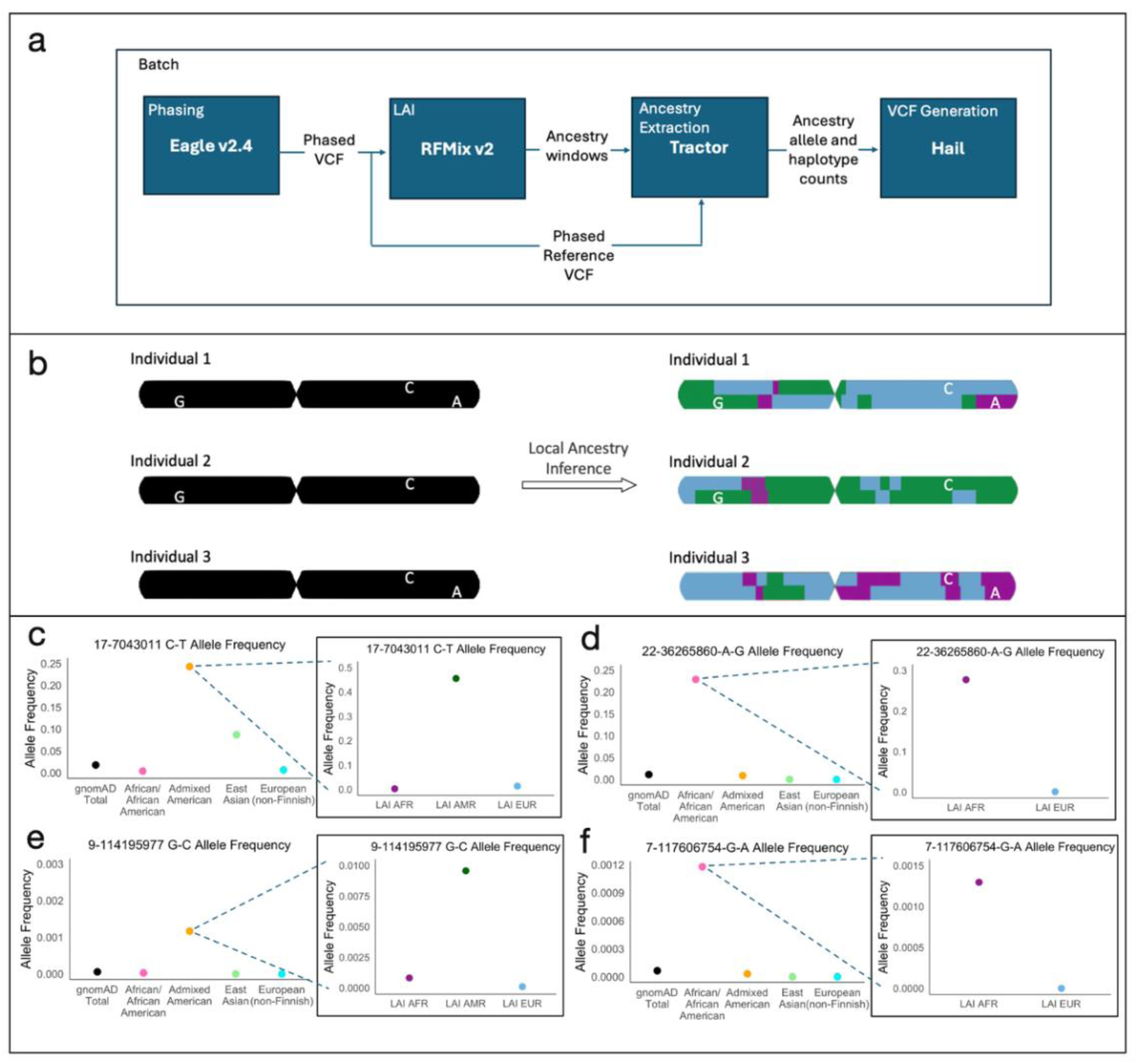
Local ancestry better resolves ancestral allele frequencies. **a.** Pipeline utilized for LAI frequency data generation**; b.** Schematic illustrating improved frequency resolution thanks to LAI. A single chromosome from three individuals is shown, with non-reference alleles at 3 SNPs (indicated by G, C, and A). After performing LAI, we observe that allele G is only found on the green ancestry, allele A only on the purple ancestry, and allele C can be found across all three; **c.** AF distribution of a *SLC16A11* variant linked to diabetes risk^11^; **d.** AF distribution of an *APOL1* variant associated with increased susceptibility to kidney disease^13^; **e.** AF distribution of a *COL27A1* variant implicated in Steel syndrome^14^; **f.** AF distribution of a *CFTR* variant linked to cystic fibrosis^15^.

LAI is useful in resolving allele frequencies for rare variants as well as common. Variants in the *COL27A1* gene have been shown to cause Steel syndrome and suggest a founder mutation effect in Puerto Rico^14^. Within gnomAD, the causal variant 9-114195977-G-C^14^ has an allele frequency of 0.1% in the Admixed American genetic ancestry group, while the gnomAD-wide allele frequency is 0.006% (**Fig. 2e**). Separating this variant’s gnomAD Admixed American genetic ancestry group frequency into continental allele frequency estimates, we see that the variant occurs almost at a 1% allele frequency in LAI-AMR regions and closer to 0% in LAI-AFR and LAI-EUR regions, reinforcing a founder effect hypothesis and again showing the usefulness of local ancestry in large genomic datasets. Another example is the gnomAD variant 7-117606754-G-A in *CFTR*, associated with cystic fibrosis^15^. This variant shows a 0.1% allele frequency in LAI-AFR regions of the African/African American genetic ancestry group, compared to a 0.007% gnomAD-wide frequency (**Fig. 2f**).

Allele frequency data is used for more than just clinical interpretation and pathogenicity. Population allele frequency (AF) databases have recently been utilized to estimate aggregate carrier frequencies and genetic prevalence, providing insights into the potential number of affected individuals^16^. LAI data enhances these estimations, enabling more precise predictions of populations with potentially higher genetic prevalence. For instance, the pathogenic variant 2-168970131-G-A (p.Arg575Ter) in the *ABCB11* gene, which causes Bile Salt Export Pump (BSEP) deficiency resulting in cholestasis^17^, has a frequency of 0.04845% in the Admixed American genetic ancestry group, the highest observed across all gnomAD genetic ancestry groups. The cumulative carrier frequency of all likely pathogenic/pathogenic (LP/P) variants in *ABCB11* across genetic ancestry groups suggests that the 2-168970131-G-A variant leads to a slightly higher carrier frequency in this group when compared to the total estimate^18^. However, analysis of LAI data revealed that this variant was exclusively observed in the LAI-EUR regions of Admixed American samples. Consequently, the elevated carrier frequency, and thus the estimated genetic prevalence, primarily pertains to individuals of higher LAI-EUR ancestry within the broader genetic ancestry inferred label. These variant examples highlight the importance of more precise allele frequencies in accurately identifying the distribution of genetic risk factors within diverse and admixed genetic ancestry groups.

### Divergence in allele frequencies across ancestry tracts

While many variants are similar in their ancestry-specific allele frequency estimates, many are also highly divergent (**Supplementary Fig. 3**). The overall correlation in ancestry-specific allele frequencies between the LAI-AFR and LAI-EUR tracts in the African/African American group was 0.83. For the Admixed American group, the correlations were 0.80 between LAI-AFR and LAI-EUR AFs, 0.73 between LAI-AFR and LAI-AMR AFs, and 0.84 between LAI-AMR and LAI-EUR AFs. Allele frequency variance was highest within the LAI-AMR tract (0.0447), compared to the LAI-AFR (0.0367) and LAI-EUR tracts (0.0356), suggesting greater heterogeneity among variants assigned to this ancestry. This elevated variance may reflect historical bottlenecks or founder effects that have shaped allele distributions in LAI-AMR haplotypes. Moreover, pairwise Levene’s tests for equality of variances between ancestry tracts were all highly significant (p < 1 × 10⁻¹⁶). Strikingly, in the African/African American genetic ancestry group, 85.1% of variants with LAI data exhibit at least a twofold greater difference between ancestry-specific AFs (**Fig. 3a**). The majority of these variants (69%) have an LAI-AFR AF that is at least two times greater than the LAI-EUR AF, while 16.1% show the opposite pattern. Similarly, in the Admixed American genetic ancestry group, 78.5% of variants demonstrate this level of divergence between ancestry-specific AFs (**Fig. 3b**). Among these, 41.6% of variants have an LAI-AFR AF that is two times greater than those of either other ancestry’s tracts, while the remaining variants show a dominant LAI-EUR AF (27.5%) or LAI-AMR AF (9.4%). Overall, LAI-AFR tracts consistently exhibit higher AFs compared to LAI-EUR or LAI-AMR tracts, both when sites that are monomorphic in the other ancestries are considered but also when excluding such loci. To complement this analysis, we also evaluated ancestry-specific depletion - variants whose AF in one ancestry tract is less than or equal to half that of the others (**Fig. 3c-d**) providing a more comprehensive genome-wide view of how allele frequencies diverge across ancestries. When restricting to monomorphic sites in the depleted tracts, 62.2% of variants in the African/African American group and 26.2% in the Admixed American group met this depletion criterion.

**Figure 3:**
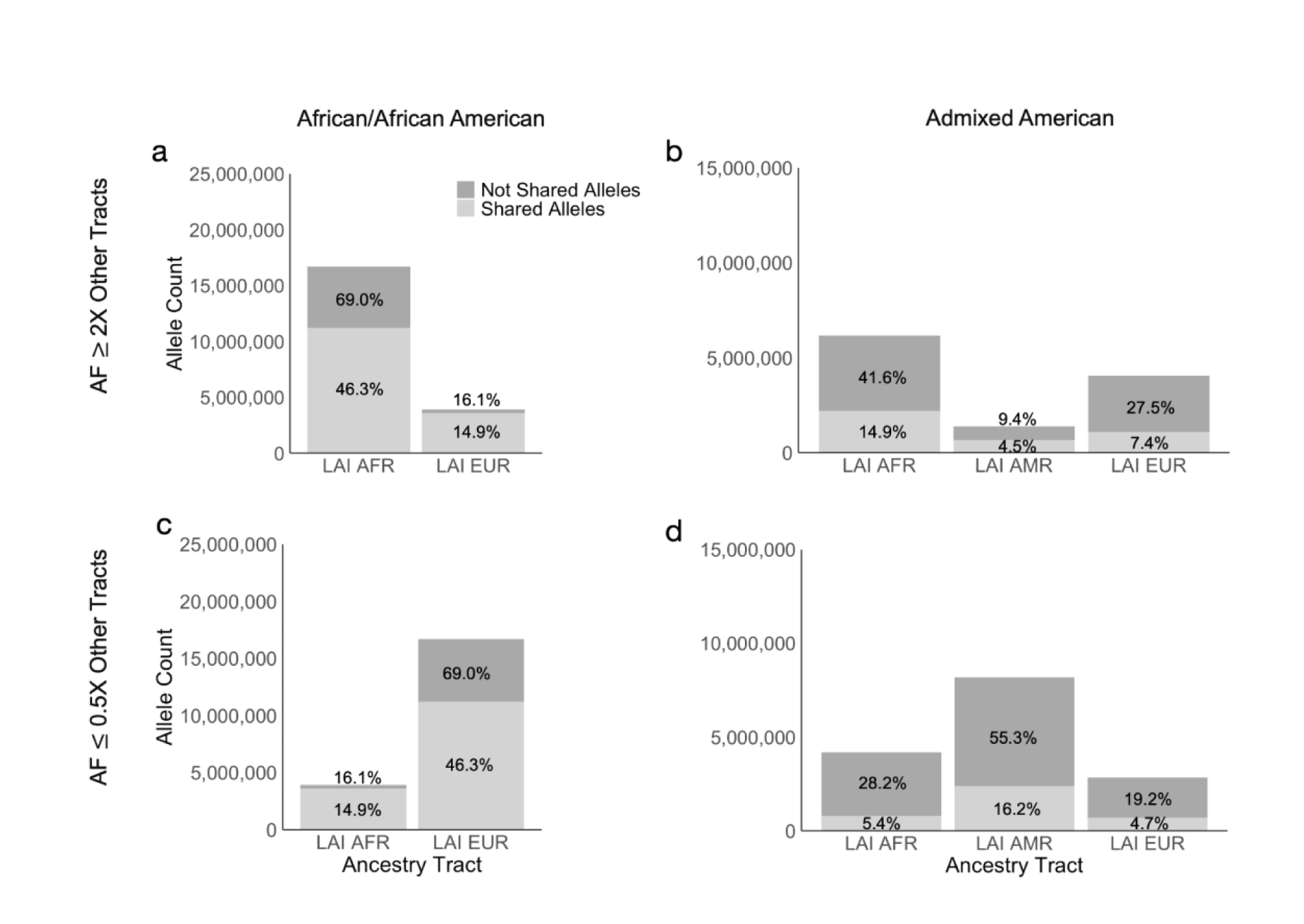
Distribution of variants with enriched or depleted ancestry tract allele frequencies across genetic ancestry groups. The total percentage for each local-ancestry background is indicated on the bars. **a,b.** Variants with ancestry-specific allele frequencies at least 2X higher than the other tracts within the African/African American and Admixed American groups, respectively. **c.d.** Variants with ancestry-specific allele frequencies at most 0.5X lower than the other tracts within the African/African American and Admixed American groups, respectively.

### Local ancestry-informed frequencies can better resolve variant clinical relevance

Visualizing the site frequency spectrum (**Fig. 4, Supplementary Fig. 4, Supplementary Table 1**) highlights the allele frequency distributions across two genetic ancestry groups classified by their ClinVar annotations^19^. Of the LAI variants, only a small fraction have corresponding ClinVar annotations: 131,886 out of 14,804,206 (0.9%) in the Admixed American and 183,164 out of 24,204,574 (0.8%) in the African/African American genetic ancestry groups. Because single-submitter and no-criteria entries may be less reliable, we applied an additional QC filter, retaining only annotations with ≥ 2 ClinVar stars, resulting in variant counts of 73,087 and 96,851, respectively. We additionally examined the number of variants with a tract-specific allele frequency at least twice as high as the other ancestral tract frequencies: the African/African American group has 75,756 variants when including all shared alleles (66,100 variants when considering only monomorphic variants), and the Admixed American group has 53,588 variants when including all shared alleles (24,683 variants when considering only monomorphic variants)(**Supplementary Fig. 5**). The results indicate that most variants across all ClinVar categories — including pathogenic/likely pathogenic, benign/likely benign, uncertain significance, and conflicting classifications of pathogenicity — are rare, with AFs concentrated between 0 and 0.1 in all genetic ancestry groups, regardless of ClinVar filtering (**Supplementary Fig. 6**). To further investigate if benign classifications may be driven primarily by frequency thresholds, we zoomed-in on benign variants and again on benign high-confidence ClinVar variants (≥2 stars) with allele frequency <0.1 and <0.01 (**Supplementary Fig. 7**). We conducted a two-sample Kolmogorov–Smirnov test on benign/likely benign ClinVar variants in the African/African American group, comparing the SFS between LAI-AFR and LAI-EUR. The test yielded a KS statistic of 0.278 with a p-value near zero (p = 10^-100), indicating a significant difference between the two distributions. The broader distribution of benign variants in African/African American groups reinforces the need to include diverse ancestry groups in genomic studies, as it demonstrates how genetic diversity impacts allele frequency distributions and, by extension, clinical variant interpretation.

**Figure 4:**
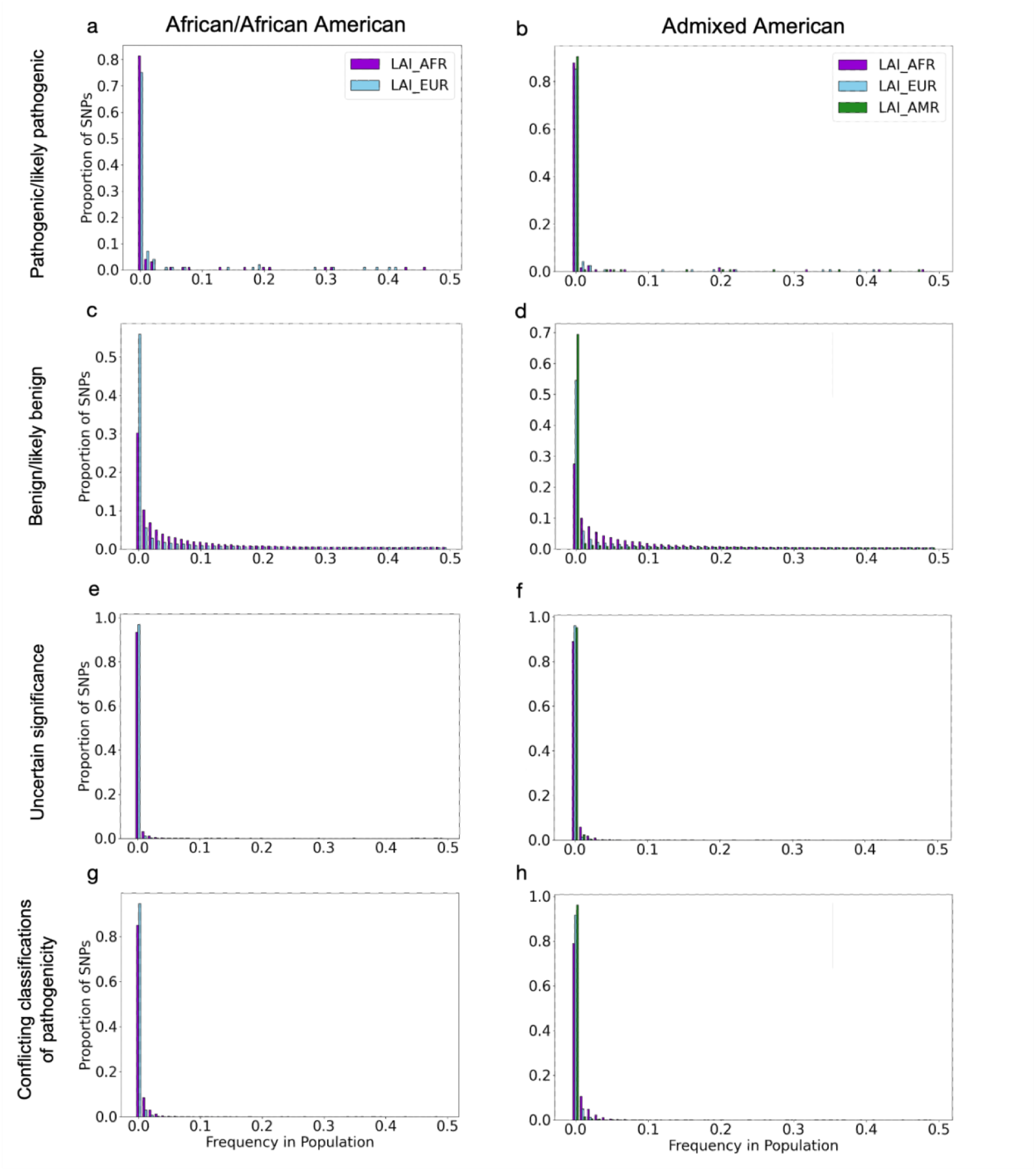
Site frequency spectrum for the inferred African/African American and Admixed American genetic ancestry groups based on ClinVar classifications. LAI-derived AFs for component ancestries are indicated with colors.

We additionally investigated the frequency of ancestry-enriched AFs exceeding those of grpmax (formerly popmax)^2^, which measures the highest allele frequency observed in any gnomAD genetic ancestry group. As grpmax is often used to inform a variant classification of benign/likely benign, better resolving allele frequencies across ancestral haplotypes with LAI provides a more precise estimate of frequencies that may change clinical interpretation. We found that across sites for both genetic ancestry groups, 22,337,592 sites exhibited a new LAI-based AF that was higher than the existing grpmax, representing 81.49% of the total 27,412,225 sites with available data. Specifically, 74.45% had tract AFs surpassing grpmax for the African/African American group, while the Admixed American group exhibited 64.95% exceeding grpmax. We also compared the tract AFs and grpmax AFs in each genetic ancestry group based on ClinVar classifications (**Supplementary Table 2)**. We found that about 87.69% and 89.32% of ClinVar variants with a higher tract AF than grpmax AF were classified as ‘Benign/Likely Benign’ in the African/African American and Admixed American genetic ancestry group, respectively.

To further explore the clinical relevance of LAI-informed allele frequencies, we focused on variants below the commonly used 5% allele frequency threshold, which is treated as stand-alone benign evidence under ACMG’s “BA1” guideline^20^ (**Supplementary Table 3**). Comparing the former grpmax AF to LAI-informed tract AFs in this critical range revealed that for the Admixed American group, 563,042 of 8,527,238 variants (∼6.6%) had a tract AF exceeding 5%. In contrast, 475,698 of 15,775,667 variants (∼3.0%) in the African/African American group showed the same pattern. To determine whether these shifts persisted among clinically curated variants, we repeated this analysis using only ClinVar variants with ≥ 2 stars ^20^. Among high-confidence variants, 3,164 of 31,485 variants (∼10.0%) in the Admixed American group and 2,684 of 42,461 variants (∼6.3%) in the African/African American group had tract AFs above 5% that exceeded their grpmax values. These findings highlight that ancestry-specific allele frequency information can better assist in the identification of likely benign variants. We additionally conducted a similar analysis stratifying by CADD PHRED score thresholds (≥10 and ≥15, corresponding to the top 10 and 5 percent of deleterious variants in the genome)^21^, comparing the frequency of variants where LAI-informed AF exceeded 5% and grpmax. For the Admixed American group, 6.1% of variants with CADD ≥10 and 5.7% with CADD ≥15 met this condition. For the African/African American group, corresponding proportions were 2.7% and 2.4%, respectively. These trends reinforce the observation that local ancestry resolution meaningfully alters the interpretation of both potentially benign and functionally consequential variants. Using grpmax AF’s without consideration of LAI could lead to inaccurate variant classification in clinical studies, particularly when evaluating the pathogenicity of variants in underrepresented populations.

In sum, we anticipate this finer resolution of allele frequencies thanks to local ancestry deconvolution will improve the interpretation of variation by contextualizing variants to their approximate geographical origin and allowing for more accurate allele frequencies to be calculated per genetic ancestry background. This improved resolution can lead to more effective identification of functionally important risk variants and a more accurate spectrum of allele frequencies. These refined estimates are freely downloadable and integrated into the widely used gnomAD browser for improved interpretation of allele frequencies across continentally defined genetic ancestries.

## DISCUSSION

An outstanding question in genomics, with far-reaching clinical and research implications, is the degree of concordance in allele frequencies across different ancestral backgrounds. Allele frequencies vary dramatically across the globe^22,23^ due to features such as demography, genetic drift, and natural selection. Deconvolving from which continental ancestry a variant is derived allows us to compute more accurate expected allele frequencies for each continental background, instead of aggregating frequencies across recently admixed individuals regardless of the ancestral origin of the variant’s haplotype^24^. The large sample size of gnomAD enables us to examine allele frequency trends on a more global scale, and with the integration of LAI, we achieve a finer resolution of ancestry-specific frequency estimates, improving the utility of the resource. In some cases, updated allele frequencies that take into account local ancestry differ significantly from their corresponding genetic ancestry group-level frequencies, and even update the highest allele frequency observed gnomAD-wide (grpmax), and/or increase frequency estimates past commonly used thresholds for estimating pathogenicity, which may impact the clinical interpretation of a variant (**Supplementary Fig. 1**).

Our analyses further reveal that a substantial number of variants exhibit highly divergent allele frequencies across ancestry backgrounds: within the African/African American and Admixed American genetic ancestry groups, 85.1% and 78.5% of variants with LAI data, respectively, show at least a twofold difference between ancestry-specific frequencies. Allele frequency differences observed across local ancestry backgrounds may arise from a combination of evolutionary and demographic forces, including genetic drift, founder effects, and natural selection. While we do not attempt to infer the specific historical or selective pressures underlying the observed patterns in this study, the extensive allele frequency divergence we report highlights the importance of modeling ancestry at finer scales. Future work leveraging these local ancestry-informed frequencies may provide insight into the history of population-enriched variants, particularly as more detailed demographic models and functional data become available.

Additionally, we found that across both genetic ancestry groups, 81.49% of the 27,412,225 sites analyzed would receive an increased grpmax value when using LAI data. When examining these variants in ClinVar, approximately 88% of those with LAI-inferred frequencies higher than grpmax were classified as ‘Benign/Likely Benign’. This suggests that many variants, correctly classified as benign using ancestry-informed data, might be underestimated or misclassified without this information. This misestimation is especially relevant near clinical decision thresholds. The ACMG guidelines designate a AF ≥5% as strong benign evidence (BA1)^25^, yet our analysis shows that many variants below this threshold by grpmax may exceed it in one or more ancestry tracts. Notably, over 10% of high-confidence ClinVar variants in the Admixed American group and 6.3% in the African/African American group cross this critical boundary when LAI is incorporated. These shifts were consistent even when stratifying by predicted functional impact using CADD scores^21^. In both genetic ancestry groups, variants with CADD ≥10 and ≥15 also exceeded the 5% threshold when tract AFs were used in place of grpmax. In terms of direct patient impact, reclassifying these variants as benign spares patients the uncertainty, repeated imaging, and potential invasive interventions they might otherwise undergo. These findings suggest that LAI enhances variant interpretation by identifying variants that, while rare globally, are common in specific ancestries—supporting their benign classification across all populations and refining our understanding of functional variant distribution across populations.

Ancestry-specific allele frequencies can additionally help refine genetic prevalence estimates for recessive diseases. Understanding the estimated proportion of a population that has a causal genotype for a genetic disorder, and whether there is a significantly higher risk in a specific genetic ancestry group or not, is a key question for understanding the global impact of a disease. Local ancestry has the potential to refine these estimates further and provide the community with a more accurate prediction of how many people have a given disease. While a 0.1% frequency cut-off is still more common than many pathogenic disease-causing variants, in the future local ancestry will be able to be applied to more rare variants, which will have an even greater effect on clinical care.

Local ancestry resolved variant allele frequencies can also help in the interpretation of GWAS results^5,26^. Differentiating between genetic ancestry group-level allele frequencies and those enriched in specific haplotypes can streamline the search for disease-associated variants with elevated importance for particular genetic ancestries^27^. For example, we demonstrate that the gnomAD variant 17-7043011-C-T in *SLC16A11*^11^, associated with type 2 diabetes risk, is observed at a 14% frequency in the Admixed American population compared to 2% gnomAD-wide. When partitioned by continental ancestry, this variant occurs at a 45% frequency in LAI-AMR regions of admixed genomes, while it is nearly absent in LAI-AFR and LAI-EUR regions. This, and the other vignettes highlighted, showcases how local ancestry inference better resolves allele frequencies in both rare and common key variants.

We recognize that genetic ancestry is inherently continuous and not discrete. For the purposes of local ancestry inference, we approximate ancestry as originating from discrete source populations, a necessary simplification for modeling recently admixed populations. Our approach does not assume that present-day populations are strictly discrete entities but rather leverages the fact that the major sources of genetic variation in recently admixed populations can be well-modeled by a small number of continental reference populations. This assumption holds well for cases of recent admixture, however, as one moves beyond recent admixture events such ancestry becomes increasingly diffuse. We additionally appreciate that continental regions are not monoliths, but contain numerous ancestries therein. Our method is not inherently limited to broad continental ancestries, and in principle, it could be extended to analyze intra-continental variation given appropriate reference data. However, the resolution of such analyses depends on the availability and comprehensiveness of reference panels as well as the degree of genetic differentiation between modeled ancestries^28^. For this reason, we are currently limited in resolving LAI to the continental rather than subcontinental level. We look forward to resolving finer-scale ancestries in the future as reference panels supporting such analyses continue to expand.

This release marks the first time local ancestry has been systematically incorporated into a globally-representative resource like gnomAD, setting a new standard for genomic inclusion. This release of local ancestry-informed frequency data is currently available for the Admixed American and African/African American genetic ancestry groups. Our computational pipeline, written in Hail Batch (see *Methods*), is publicly available via GitHub. The allele frequencies and quality metrics can be downloaded or interactively explored on the gnomAD browser (https://gnomad.broadinstitute.org/). We hope this data can be used to enhance the scientific community’s ability to clinically interpret variation for those of all genomic backgrounds.

## METHODS

### Sample selection and quality control

Genetic ancestry group labels were assigned to samples as previously described^1^ - briefly, principal components (PCs) computed from hail’s hwe_normalized_pca^2^ were used as features when training a random forest classifier using Thousand Genomes Project/Human Genome Diversity Sample reference samples with known continental origin (**Fig. 1a**). A continental genetic ancestry group was assigned to all samples for which the probability of that continental genetic ancestry group was > 90% according to the random forest model. Samples were subset based on inferred genetic ancestry groups, limiting to bi-allelic variants with low genotype missingness (i.e., call rate > 0.9) and an allele frequency greater than 0.1% within the Admixed American and African/African American gnomAD genetic ancestry groups. Additionally, an allele count (AC) threshold of AC > 7 was applied for each genetic ancestry group. The 7,612 Admixed American and 20,250 African/African American whole genome sequencing samples from gnomAD v3.1 used here include all individuals in the database except those from the 1000 Genomes Project (1kGP) and Human Genome Diversity Project (HGDP) joint callset^3^, which were used to curate reference panels for LAI in each genetic ancestry group.

### Global ancestry deconvolution

To explore the overall population composition of the gnomAD African/African American and Admixed American samples, we conducted principal components analysis (PCA) and unsupervised ADMIXTURE^4^ to infer global genetic ancestry proportions. Informed by these global ancestry estimates, we set k=3 for the Admixed American genetic ancestry group (including all reference samples from 1kG/HGDP labeled as belonging to the AFR, EUR, and AMR superpopulations). For the African/African American genetic ancestry group, we set k=2 (using just continental reference samples from the AFR and EUR superpopulations). In this work, we focus on two admixed genetic ancestry groups represented in gnomAD: the African/African American and Admixed American groups. The underlying composition of the genomes of individuals from these groups contain complex mosaics of ancestries shaped by recent admixture events involving African, European, and Indigenous American sources^5–7^. African/African American individuals typically carry a majority of West African ancestry, with variable proportions of European and, to a lesser extent, Indigenous American ancestry. Admixed American individuals generally derive their genomes from a mixture of Indigenous American, European (primarily Iberian), and African ancestries, with substantial regional and individual-level variation. These groups have historically been underrepresented in genomic studies, and their complex ancestry patterns make them particularly well-suited for local ancestry-informed analyses.

### Local ancestry inference pipeline

To infer local ancestry data in the gnomAD Admixed American and African/African American group, we built an open-source LAI pipeline. **Fig. 2a** and **2b** shows a schematic of the software pipeline, which includes four primary steps: phasing, LAI, ancestry extraction/dosage calculation, and LAI-informed allele frequency generation. We used Eagle^8^ for separately phasing the Admixed American reference panel and for joint phasing of the African/African American group, as it is computationally tractable at scale^8^. We used RFmixV2^9^ for LAI, which requires a reference panel representative of the continental ancestries present within the samples. Our reference panel was composed of the harmonized HGDP and 1kGP3 “AMR” samples to capture the Amerindigenous genetic components in the Admixed American group, “EUR” samples for the European components, and “AFR” samples to capture continental African genetic ancestries. RFmixV2^9^ was run with one EM iteration for the Admixed American samples to account for the high amount of admixture present in the three-way reference. For the curated reference for the African/African American group, we excluded the Americans of African Ancestry in Southwest USA (ASW) and African Caribbeans in Barbados (ACB) due to their higher level of admixture. Given the substantially larger sample size of the African/African American genetic ancestry group in gnomAD, reducing admixture within the reference panel and therefore removing the need for an EM iteration dramatically reduces the required computing time.

To extract ancestral segments and calculate ancestry-specific minor allele dosages, we used the tool Tractor^10^. Due to computational constraints, samples from the inferred African/African American genetic ancestry group were run in batches of 6,750 samples for chromosomes 1 and 3-12, and batches of 3,375 samples for chromosome 2 (**Supplementary Table 4**) for LAI. Since the same reference panel was used across batches, this is not expected to impact results. Finally, we tabulated local ancestry-informed allele frequencies using a custom Python script that utilizes the Hail library^2^. To ensure consistency with the most up-to-date dataset available to users, we report total allele frequencies and genetic ancestry group AFs from gnomAD v4.1.0 for all example variants. While LAI calls remain unchanged between gnomAD versions, the aggregate allele frequencies (both total and stratified by genetic ancestry group) have been updated in v4.1.0 due to the inclusion of additional diverse samples. To evaluate the pipeline’s sensitivity, we applied it to a simulated admixed American truth dataset and computed the true positive rates of LAI^11^ (**Supplementary Fig. 8**). We also validated our pipeline by comparing global ancestry proportions estimated by ADMIXTURE to those from our LAI pipeline results (**Supplementary Fig. 9**). The entire pipeline is unified using the Hail Batch Python module for scalability and is available at [github.com/broadinstitute/gnomad_local_ancestry].

## DATA AVAILABILITY

Data can be downloaded from or browsed interactively at [gnomad.broadinstitute.org].

## CODE AVAILABILITY

Code is freely available at [github.com/broadinstitute/gnomad_local_ancestry], with a detailed README outlining the LAI pipeline and step-by-step instructions for running it.

## ACKNOWLEDGMENTS

We wish to thank Alicia Byrne, Anna Lewis, and Alham Saadat for literature review; Jackie Goldstein, Dan King, and Ben Weisburd for pipeline scalability assistance; Marcos Santoro for providing LAI reference panel guidance; Julia Goodrich for pipeline code review. This work was funded by the National Institutes of Health (K01 MH121659 and R01 HG012869 to E.G.A.; P.K. was supported by T32 GM139534-03 and T32 GM136554-05).

## AUTHOR CONTRIBUTIONS

P.K. and M.W. led data analysis with support from G.T., K.C., S.M.B., J.H.M., H.L.R., M.J.D. and K.J.K. advised on analyses. P.D. and N.W incorporated LAI data in the browser. E.G.A. designed the study and supervised analysis. All authors have read and approve this manuscript.

