## Supplementary document for "Improved Allele Frequencies in gnomAD through Local Ancestry Inference"

### SUPPLEMENTARY FIGURES

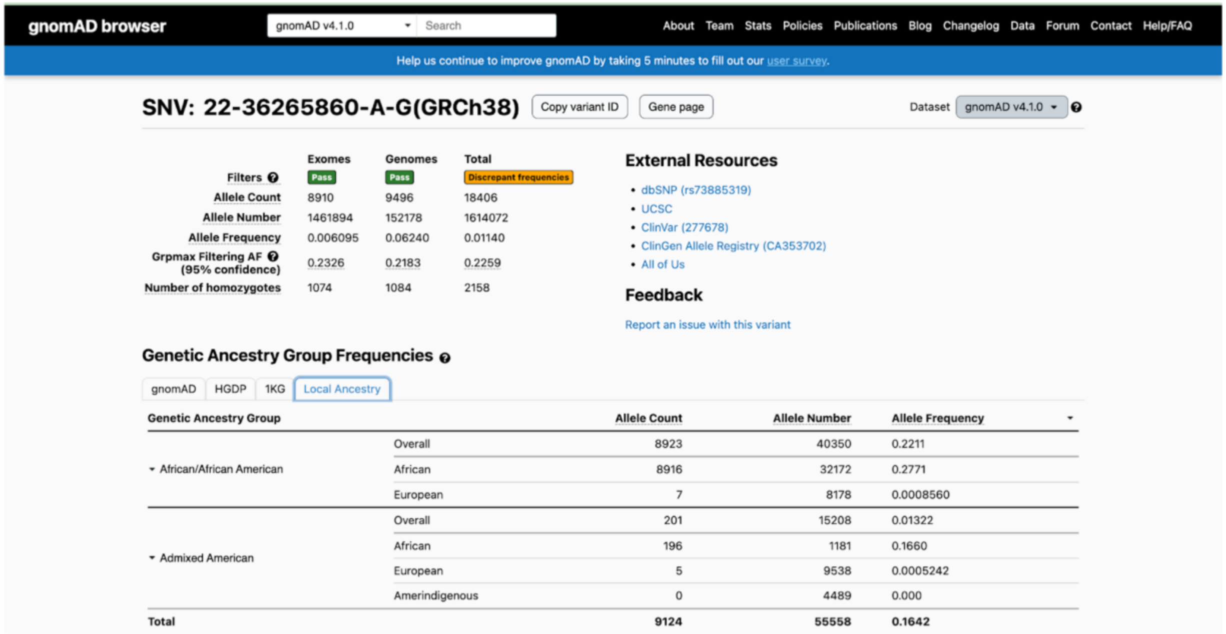

**Supplementary Figure 1. Screenshot of local ancestry resolved allele frequencies in the gnomAD browser.** To access the local ancestry data for a variant within the browser, select the Local Ancestry tab within the population frequency table on the variant's page. If local ancestry data is available for the variant, the ancestral allele count (AC), allele number (AN), and allele frequency (AF) estimates will be displayed.

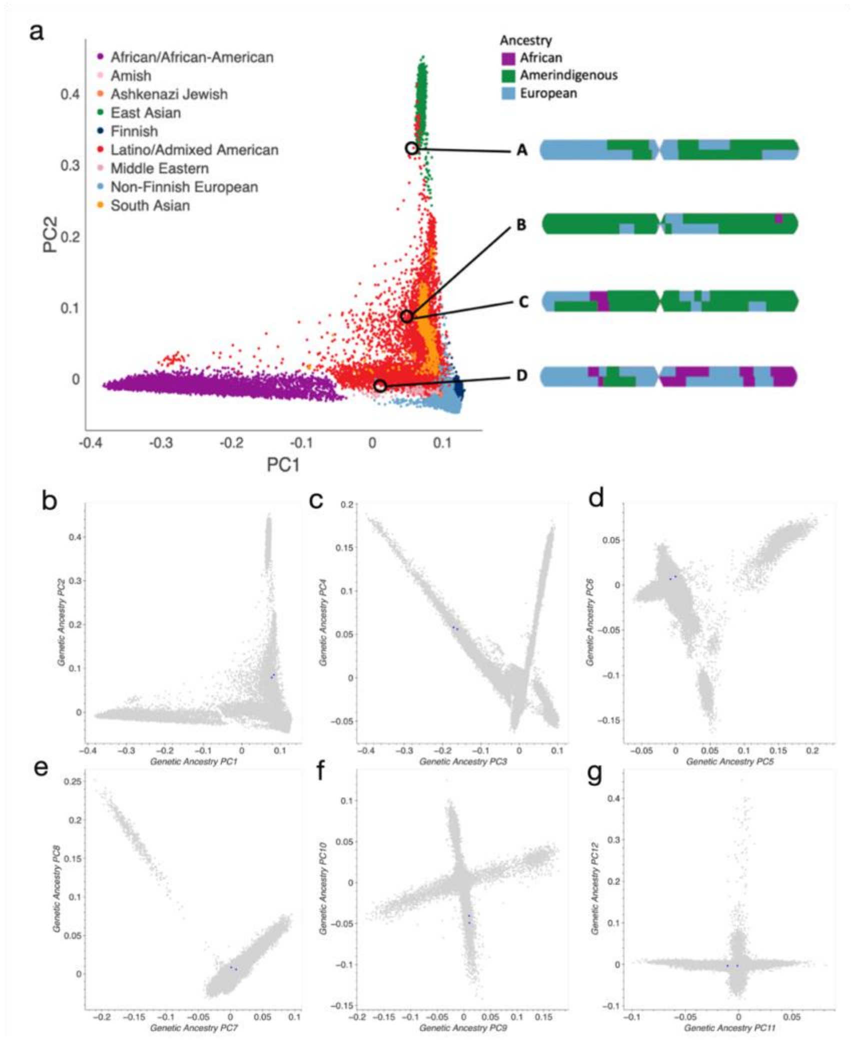

**Supplementary Figure 2: Population structure of gnomAD v3.1 samples. a.** Principal component analysis of all samples in gnomAD v3.1 projected onto the first two principal components, colored and labeled according to the flagship gnomAD genetic ancestry scheme and naming convention at the time. Four example Latino/Admixed American individuals (A–D) are highlighted to illustrate the heterogeneity in global and local ancestry within a single inferred genetic ancestry group. **b-g.** Two samples (B, C) are additionally featured throughout deeper PCs to demonstrate that even samples with similar global PC coordinates have distinct local ancestry compositions. Note that individuals B and C (highlighted in blue) remain in close proximity across higher PCs, despite having different local ancestry configurations.

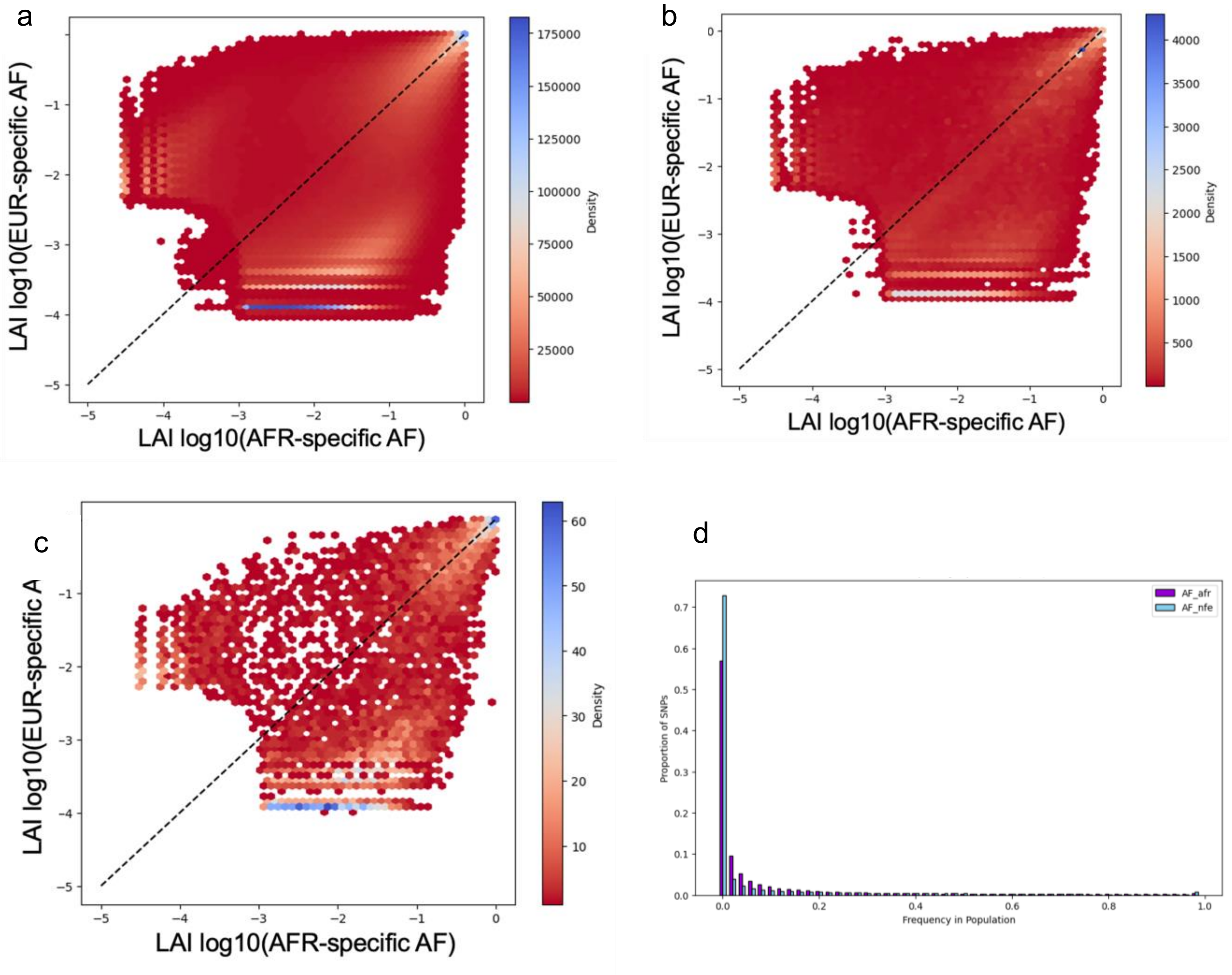

**Supplementary Figure 3: Ancestry-specific allele frequency spectra in the African/African American genetic ancestry group.** The black dotted line represents a 1:1 ratio and the color gradient indicates density of the data. **a.** Hexbin plot of ancestry-specific allele frequencies ($\log_{10}$ AF) for all variants tested. **b.** Hexbin plot of ancestry-specific allele frequencies ($\log_{10}$ AF) for only chromosome 22, for improved visibility. **c.** Hexbin plot of ancestry-specific allele frequencies ($\log_{10}$ AF) for 10,000 random variants, for improved visibility. **d.** Site Frequency Spectrum (SRS) for the inferred African/African American genetic ancestry group.

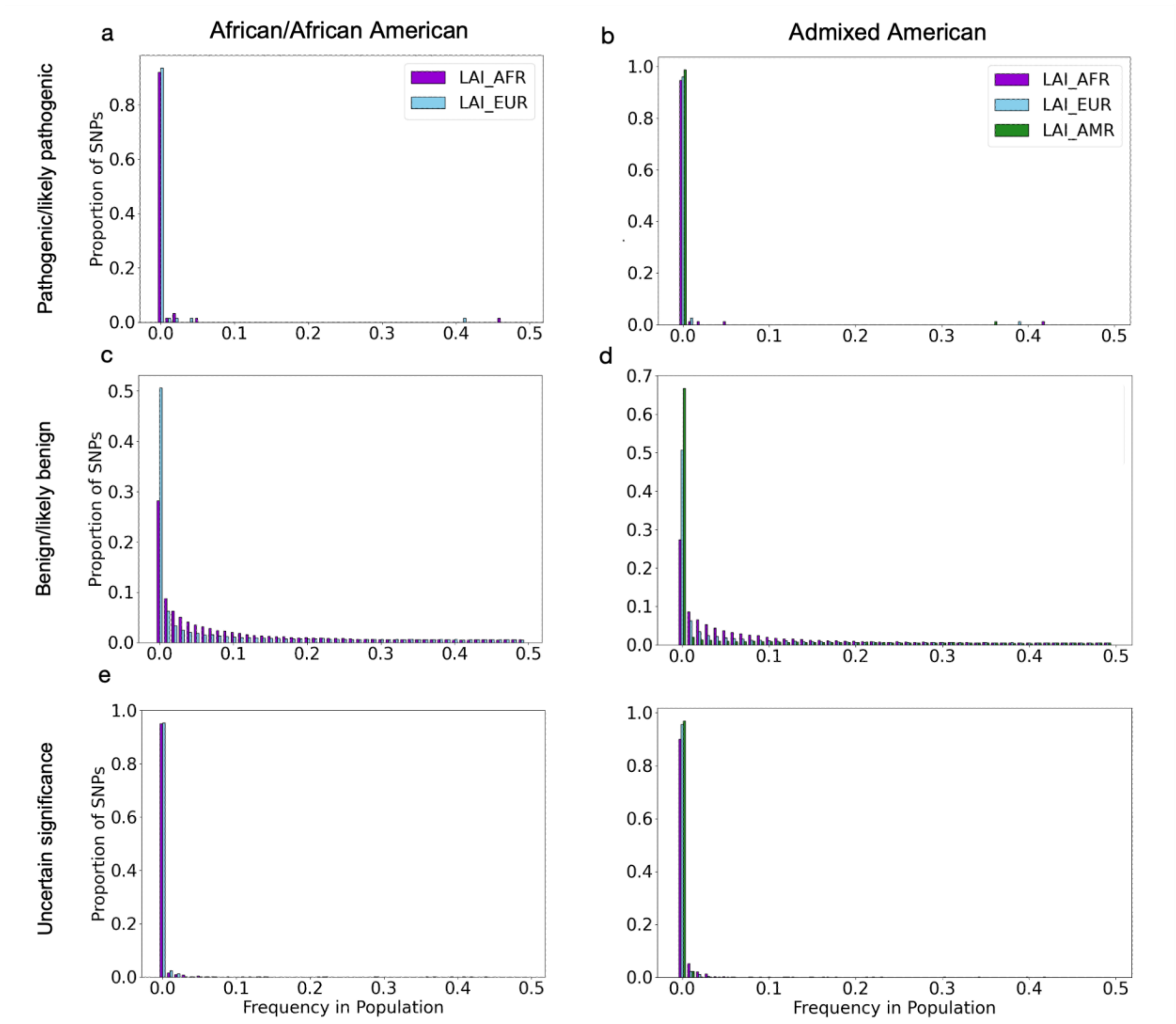

**Supplementary Figure 4: Site frequency spectrum for inferred genetic ancestry groups based on ClinVar classifications with high-confidence ClinVar variants (≥2 stars)**. LAI-derived AFs for component ancestries are indicated with colors.

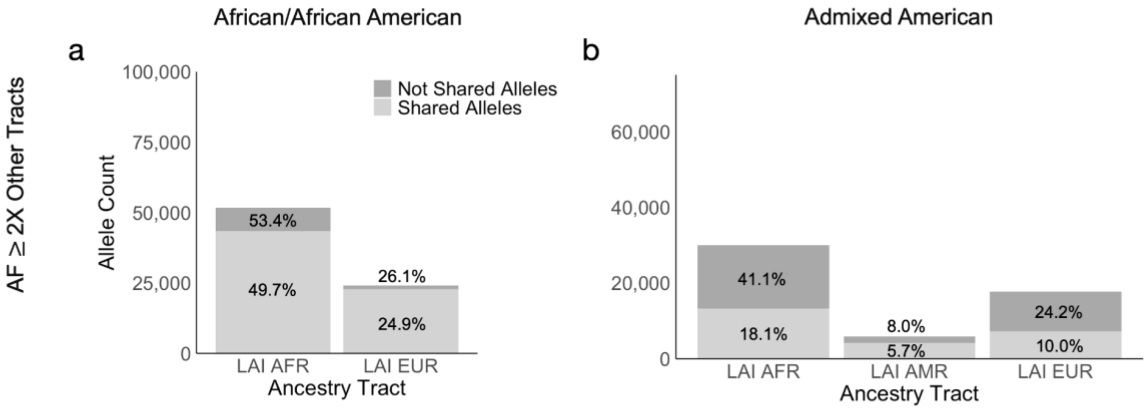

**Supplementary Figure 5: Distribution of high-confidence ClinVar variants (≥2 stars) with enriched specific ancestry tract allele frequencies ≥ 2x other ancestral tracts across genetic ancestry groups.** The total percentage for each local-ancestry informed tract is indicated on the bars. **a,b.** Variants with ancestry-specific allele frequencies at least 2X higher than the other tracts within the African/African American and Admixed American groups, respectively.

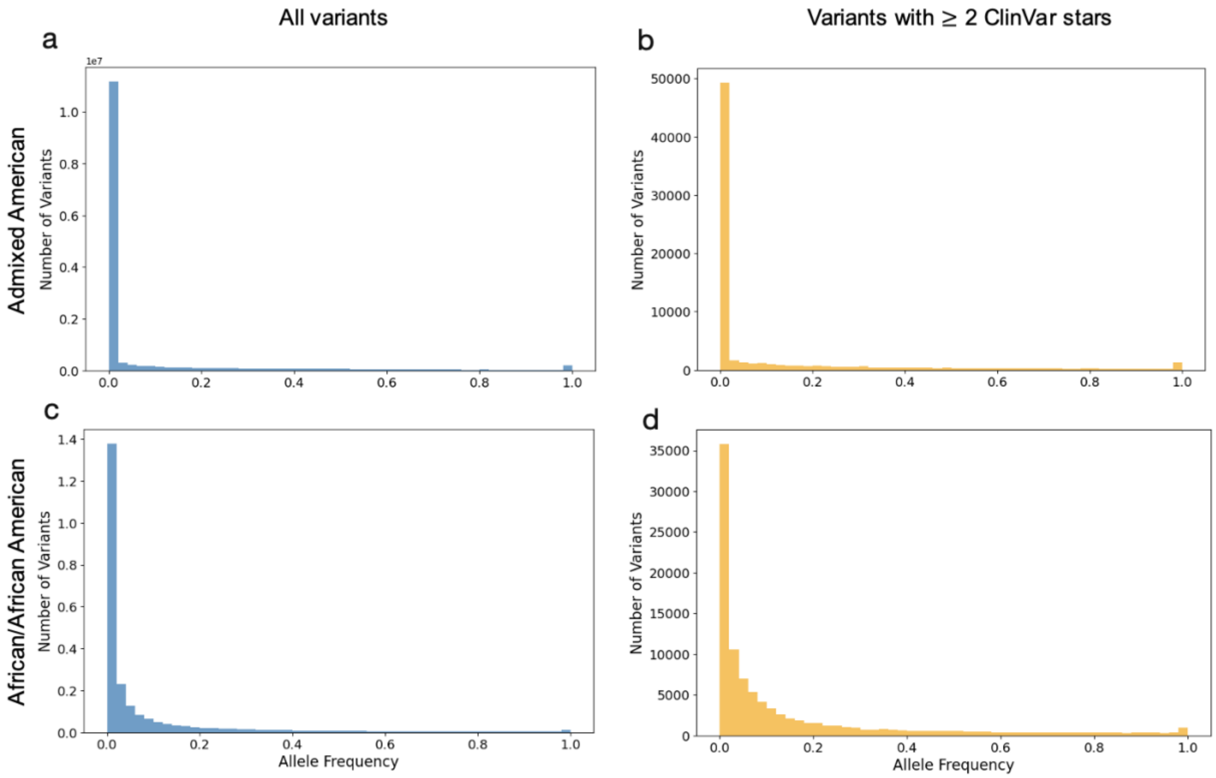

**Supplementary Figure 6: Distribution of allele frequencies in Admixed American and African/African American populations. a, c.** Site frequency spectra (SFS) of all variants with LAI-informed allele frequencies in the Admixed American and African/African American genetic ancestry groups, respectively. **b,d.** SFS of high-confidence variants (≥ 2 ClinVar stars) in the same respective groups.

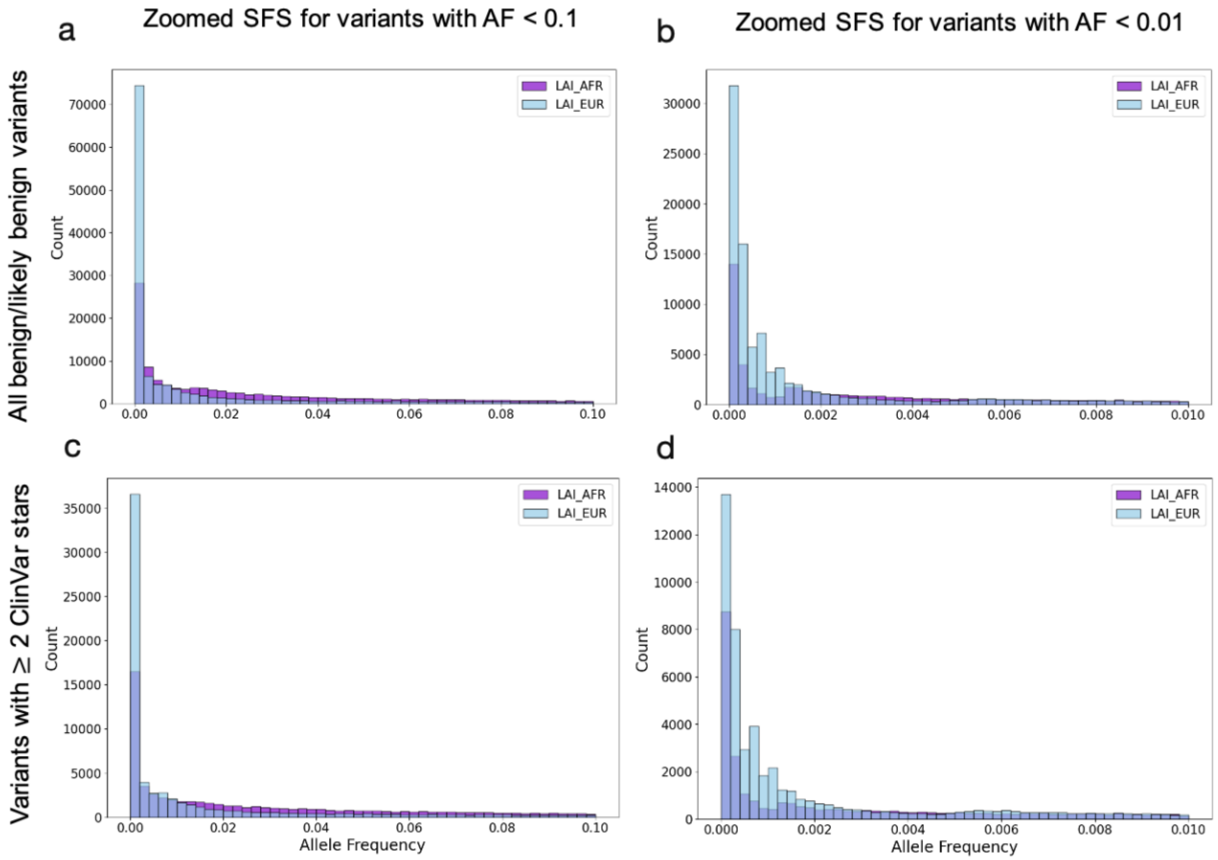

**Supplementary Figure 7: Zoomed-in site frequency spectra for benign/likely benign variants in the African/African American genetic ancestry group.** Purple and blue bars represent local ancestry-informed allele frequencies from African (LAI_AFR) and European (LAI_EUR) ancestry tracts, respectively. **a,b.** All ClinVar variants annotated as Benign/Likely Benign with AF < 0.1 and AF < 0.01. **c,d.** High-confidence Benign/likely benign variants (≥ 2 ClinVar stars) with AF < 0.1 (c) and AF < 0.01.

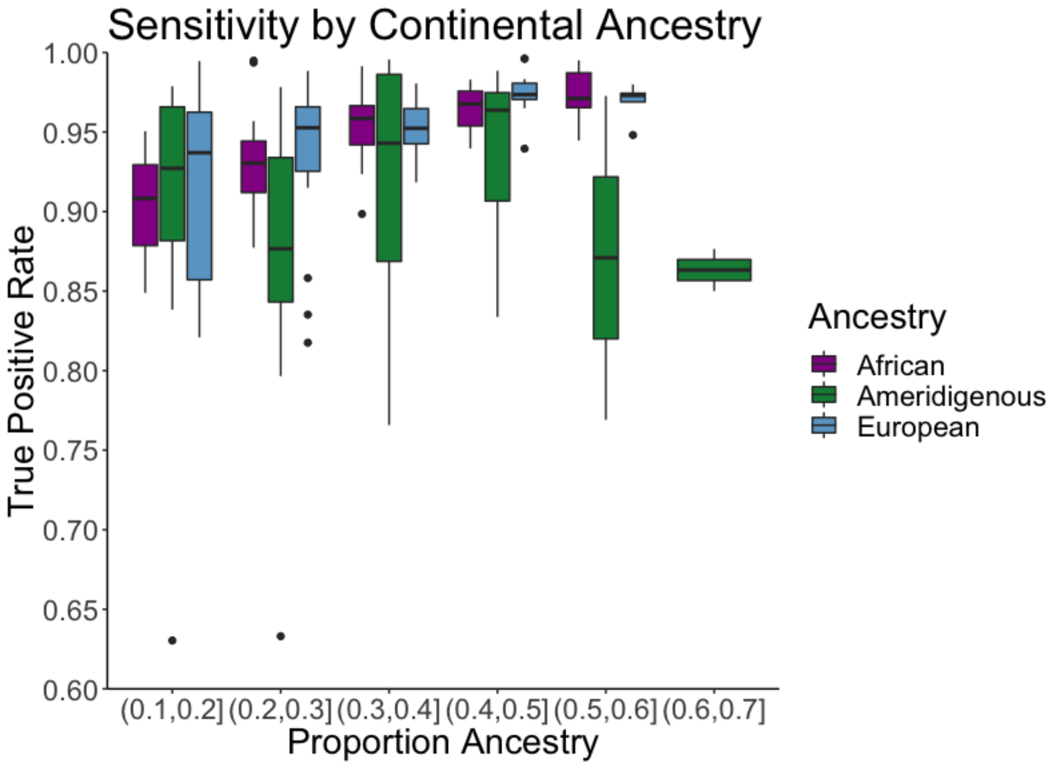

**Supplementary Figure 8: Accuracy of local ancestry estimation within ancestral backgrounds of gnomAD Latino/Admixed Americans.** The true positive rate, ascertained from conducting LAI on simulated truth data, as shown across the three modeled ancestral backgrounds. Samples were binned according to their overall global ancestry.

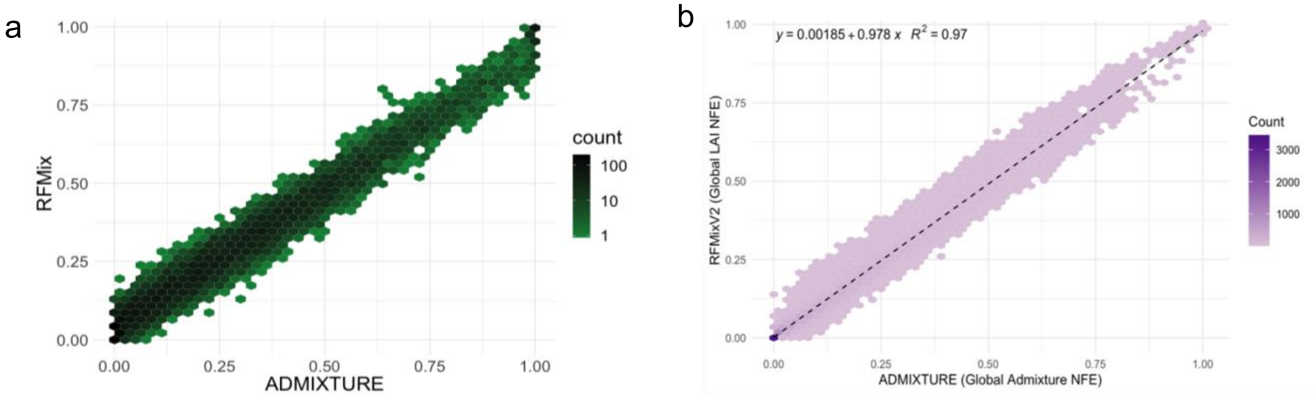

**Supplementary Figure 9: Comparison of global ancestry estimates.** Both panels show the global proportion of AFR estimated by ADMIXTURE on the x axis versus that from RFmix on the y axis. The color gradient indicates density of the data. **a.** gnomAD Admixed American genetic ancestry group. **b.** gnomAD African/African American genetic ancestry group.

**Supplementary Table 1. Number of LAI variants with ClinVar annotations by significance category and genetic ancestry group, before and after applying a high-confidence (≥2 stars) filter.**

| Genetic Ancestry Group | ClinVar Significance | Variant Count | High-Confidence (≥2 ClinVar stars) Variant Count |
|---|---|---|---|
| **Admixed American** | Benign/Likely_benign | 121,056 | 71,736 |
| | Conflicting_classifications_of_pathogenicity | 4,750 | NA |
| | Pathogenic/Likely_pathogenic | 116 | 76 |
| | Uncertain_significance | 5,964 | 1,275 |
| | Total | **131,886** | **73,087** |
| **African/African American** | Benign/Likely_benign | 166,629 | 95,238 |
| | Conflicting_classifications_of_pathogenicity | 8,062 | NA |
| | Pathogenic/Likely_pathogenic | 97 | 62 |
| | Uncertain_significance | 8,376 | 1,551 |
| | Total | **183,164** | **96,851** |

**Supplementary Table 2. ClinVar classification breakdown of variants with a change in grpmax value due to a higher local ancestry-informed allele frequency.**

| Genetic Ancestry Group | ClinVar Annotation | Allele Count | Percentage |
|---|---|---|---|
| **African/African American** | Pathogenic/Likely Pathogenic | 63 | 0.05% |
| | Benign/Likely Benign | 108,121 | 87.69% |
| | Uncertain Significance | 7,927 | 6.43% |
| | Conflicting Classifications of Pathogenicity | 7,177 | 5.82% |
| **Admixed American** | Pathogenic/Likely Pathogenic | 90 | 1.08% |
| | Benign/Likely Benign | 74,436 | 89.32% |
| | Uncertain Significance | 5,372 | 6.45% |
| | Conflicting Classifications of Pathogenicity | 3,438 | 4.13% |

**Supplementary Table 3. Number and percentage of variants where local ancestry-informed allele frequency exceeds grpmax, stratified by grpmax AF < 5% and additional variant prioritization filters.**

| | Admixed American | African/African American |
|---|---|---|
| **All variants with grpmax AF < 5%** | 563,042 variants (6.6%) | 475,698 variants (3.0%) |
| **CADD PHRED score >=10** | 32,267 variants (6.1%) | 24,336 variants (2.7%) |
| **CADD PHRED score >=15** | 10,279 variants (5.7%) | 7,394 variants (2.4%) |
| **2+ ClinVar Stars** | 3,164 variants (10%) | 2,684 variants (6.3%) |

**Supplementary Table 4. Breakdown of runtime and computational resources used for steps in the gnomAD LAI pipeline.**

| Chr | Step | Duration | Cost | Specified Machine | Split |
|---|---|---|---|---|---|
| 1 | Eagle | 2 days 19 hrs | $168.84 | n1-highmem-32 | |
| 1 | Split vcf (cohort) | 0 days 19 hrs 9 mins | $25.32 | | |
| 1 | Split vcf (reference) | 0 days 5 hrs 29 mins | $7.26 | | |
| 1 | RFMix | 0 days 6 hrs 57 mins | $17.56 | n1-highmem-32 | 3x |
| 1 | Tractor | 2 days 7 hrs | $74.97 | | 3x |
| 1 | Generate Final VCF | 0 days 6 hrs 21 mins | $8.23 | | |
| 2 | Eagle | 3 days 6 hrs | $195.71 | n1-highmem-32 | |
| 2 | Split vcf (cohort) | 0 days 7 hrs 38 mins | $9.30 | | 6x |
| 2 | Split vcf (reference) | 0 days 5 hrs 53 mins | $7.79 | | |
| 2 | RFMix | 0 days 5 hrs 31 mins | $13.98 | | 6x |
| 2 | Tractor | 1 day 6 hrs | $47.34 | n1-highmem-32 | 6x |
| 2 | Generate Final VCF | 0 days 7 hrs 9 mins | $9.24 | | |
| 3 | Eagle | 4 days 14 hrs | $145.94 | | |
| 3 | Split vcf (cohort) | 0 days 16 hrs 34 mins | $40.22 | | |
| 3 | Split vcf (reference) | 0 days 4 hrs 54 mins | $6.49 | | |
| 3 | RFMix | 0 days 6 hrs 1 min | $15.24 | n1-highmem-32 | 3x |
| 3 | Tractor | 2 days 1 hr | $67.61 | | 3x |
| 3 | Generate Final VCF | 0 days 5 hrs 49 mins | $7.52 | | |
| 4 | Eagle | 4 days 10 hrs | $141.27 | | |
| 4 | Split vcf (cohort) | 0 days 17 hrs 55 mins | $23.70 | | |
| 4 | Split vcf (reference) | 0 days 4 hrs 46 mins | $6.31 | | |
| 4 | RFMix | 0 days 6 hrs 24 mins | $16.19 | n1-highmem-32 | 3x |
| 4 | Tractor | 1 day 23 hrs | $64.62 | | 3x |
| 4 | Generate Final VCF | 0 days 6 hrs 9 mins | $7.96 | | |
| 5 | Eagle | 4 days 1 hr | $126.28 | | |
| 5 | Split vcf (cohort) | 0 days 17 hrs 16 mins | $22.46 | | |

| | | | | | |
|---|---|---|---|---|---|
| 5 | Split vcf (reference) | 0 days 4 hrs 31 mins | $5.88 | | |
| 5 | RFMix | 0 days 6 hrs 23 mins | $16.18 | n1-highmem-32 | 3x |
| 5 | Tractor | 1 day 23 hrs | $63.75 | | 3x |
| 5 | Generate Final VCF | 0 days 5 hrs 56 mins | $7.68 | | |
| 6 | Eagle | 4 days 5 hrs | $132.27 | | |
| 6 | Split vcf (cohort) | 0 days 16 hrs 35 mins | $21.56 | | |
| 6 | Split vcf (reference) | 0 days 4 hrs 14 mins | $5.51 | | |
| 6 | RFMix | 0 days 5 hrs 30 mins | $14.26 | n1-highmem-32 | 3x |
| 6 | Tractor | 1 day 21 hrs | $61.91 | | 3x |
| 6 | Generate Final VCF | 0 days 5 hrs 41 mins | $7.36 | | |
| 7 | Eagle | 3 days 20 hrs | $120.33 | | |
| 7 | Split vcf (cohort) | 0 days 15 hrs 26 mins | $20.08 | | |
| 7 | Split vcf (reference) | 0 days 4 hrs 7 mins | $5.37 | | |
| 7 | RFMix | 0 days 5 hrs 14 mins | $13.28 | n1-highmem-32 | 3x |
| 7 | Tractor | 1 day 18 hrs | $57.51 | | 3x |
| 7 | Generate Final VCF | 0 days 4 hrs 52 mins | $6.29 | | |
| 8 | Eagle | 3 days 9 hrs | $106.32 | | |
| 8 | Split vcf (cohort) | 0 days 14 hrs 48 mins | $19.25 | | |
| 8 | Split vcf (reference) | 0 days 4 hrs 9 mins | $5.41 | | |
| 8 | RFMix | 0 days 5 hrs 15 mins | $13.30 | n1-highmem-32 | 3x |
| 8 | Tractor | 1 day 16 hours | $54.91 | | 3x |
| 8 | Generate Final VCF | 0 days 4 hrs 35 mins | $5.92 | | |
| 9 | Eagle | 2 days 22 hrs | $91.57 | | |
| 9 | Split vcf (cohort) | 0 days 11 hrs 52 mins | $15.44 | | |
| 9 | Split vcf (reference) | 0 days 3 hrs 8 mins | $4.09 | | |
| 9 | RFMix | 0 days 4 hrs 12 mins | $10.66 | n1-highmem-32 | 3x |
| 9 | Tractor | 1 day 11 hrs | $48.67 | | 3x |
| 9 | Generate Final VCF | 0 days 4 hrs 9 mins | $5.36 | | |
| 10 | Eagle | 3 days 1 hr | $95.25 | | |
| 10 | Split vcf (cohort) | 0 days 12 hrs 15 mins | $15.94 | | |
| 10 | Split vcf (reference) | 0 days 3 hrs 23 mins | $4.42 | | |

| | | | | | |
|---|---|---|---|---|---|
| 10 | RFMix | 0 days 4 hrs 24 mins | $11.15 | n1-highmem-32 | 3x |
| 10 | Tractor | 1 day 6 hrs | $41.73 | | 3x |
| 10 | Generate Final VCF | 0 days 4 hrs 31 mins | $6.05 | | |
| 11 | Eagle | 3 days 2 hrs | $96.62 | | |
| 11 | Split vcf (cohort) | 0 days 12 hrs 40 mins | $16.48 | | |
| 11 | Split vcf (reference) | 0 days 3 hrs 24 mins | $4.44 | | |
| 11 | RFMix | 0 days 4 hrs 39 mins | $11.82 | n1-highmem-32 | 3x |
| 11 | Tractor | 1 day 11 hrs | $48.02 | | 3x |
| 11 | Generate Final VCF | 0 days 4 hrs 14 mins | $5.51 | | |
| 12 | Eagle | 2 days 18 hrs | $162.97 | | |
| 12 | Split vcf (cohort) | 0 days 11 hrs 41 mins | $15.21 | | |
| 12 | Split vcf (reference) | 0 days 3 hrs 3 mins | $3.98 | | |
| 12 | RFMix | 0 days 4 hrs 14 mins | $11.21 | n1-highmem-32 | 3x |
| 12 | Tractor | 1 day 9 hrs | $45.35 | | 3x |
| 12 | Generate Final VCF | 0 days 4 hrs 19 mins | $5.63 | | |
| 13 | Eagle | 2 days 2 hrs | $64.59 | | |
| 13 | Split vcf (cohort) | 0 days 8 hrs 27 mins | $11.02 | | |
| 13 | Split vcf (reference) | 0 days 2 hrs 4 mins | $3.28 | | |
| 13 | RFMix | 0 days 7 hrs 53 mins | $19.38 | n1-highmem-32 | |
| 13 | Tractor | 6 days 4 hrs | $200.16 | | |
| 13 | Generate Final VCF | 0 days 3 hrs 23 mins | $4.42 | | |
| 14 | Eagle | 1 day 23 hrs | $60.76 | | |
| 14 | Split vcf (cohort) | 0 days 8 hrs 6 mins | $10.37 | | |
| 14 | Split vcf (reference) | 0 days 2 hrs 2 mins | $2.62 | | |
| 14 | RFMix | 0 days 7 hrs 14 mins | $17.81 | n1-highmem-32 | |
| 14 | Tractor | 5 days 23 hrs | $186.85 | | |
| 14 | Generate Final VCF | 0 days 2 hrs 53 mins | $3.75 | | |
| 15 | Eagle | 1 day 17 hrs | $52.41 | | |
| 15 | Split vcf (cohort) | 0 days 6 hrs 43 mins | $8.61 | | |
| 15 | Split vcf (reference) | 0 days 2 hrs 4 mins | $2.65 | | |
| 15 | RFMix | 0 days 6 hrs 39 mins | $16.48 | n1-highmem-32 | |
| 15 | Tractor | 5 days 10 hrs | $166.56 | | |

| | | | | |
|---|---|---|---|---|
| 15 | Generate Final VCF | 0 days 2 hrs 38 mins | $3.45 | |
| 16 | Eagle | 1 day 21 hrs | $58.35 | |
| 16 | Split vcf (cohort) | 0 days 8 hrs 11 mins | $10.47 | |
| 16 | Split vcf (reference) | 0 days 2 hrs 2 mins | $2.61 | |
| 16 | RFMix | 0 days 6 hrs 51 mins | $16.87 | n1-highmem-32 |
| 16 | Tractor | 5 days 20 hrs | $182.67 | |
| 16 | Generate Final VCF | 0 days 3 hrs 1 min | $3.93 | |
| 17 | Eagle | 1 day 19 hrs | $56.01 | |
| 17 | Split vcf (cohort) | 0 days 7 hrs 24 mins | $9.47 | |
| 17 | Split vcf (reference) | 0 days 1 hr 55 mins | $2.46 | |
| 17 | RFMix | 0 days 6 hrs 29 mins | $15.98 | n1-highmem-32 |
| 17 | Tractor | 5 days 4 hrs | $190.00 | |
| 17 | Generate Final VCF | 0 days 2 hrs 31 mins | $3.28 | |
| 18 | Eagle | 1 day 17 hrs | $86.30 | |
| 18 | Split vcf (cohort) | 0 days 7 hrs 13 mins | $9.23 | |
| 18 | Split vcf (reference) | 0 days 1 hr 56 mins | $2.49 | |
| 18 | RFMix | 0 days 6 hrs 31 mins | $16.05 | n1-highmem-32 |
| 18 | Tractor | 5 days 4 hrs | $187.93 | |
| 18 | Generate Final VCF | 0 days 2 hrs 34 mins | $3.36 | |
| 19 | Eagle | 1 day 11 hrs | $44.24 | |
| 19 | Split vcf (cohort) | 0 days 5 hrs 31 mins | $6.94 | |
| 19 | Split vcf (reference) | 0 days 1 hr 31 mins | $1.90 | |
| 19 | RFMix | 0 days 4 hrs 50 mins | $12.03 | n1-highmem-32 |
| 19 | Tractor | 4 days 8 hrs | $133.14 | |
| 19 | Generate Final VCF | 0 days 2 hrs 3 mins | $2.67 | |
| 20 | Eagle | 1 day 10 hrs | $43.67 | |
| 20 | Split vcf (cohort) | 0 days 5 hrs 34 mins | $7.01 | |
| 20 | Split vcf (reference) | 0 days 1 hr 30 mins | $1.89 | |
| 20 | RFMix | 0 days 5 hrs 10 mins | $12.78 | n1-highmem-32 |
| 20 | Tractor | 4 days 10 hrs | $136.29 | |
| 20 | Generate Final VCF | 0 days 2 hrs 5 mins | $2.71 | |

| | | | |
|---|---|---|---|
| 21 | Eagle | 1 day 1 hr | $32.14 |
| 21 | Split vcf (cohort) | 0 days 3 hrs 56 mins | $4.95 |
| 21 | Split vcf (reference) | 0 days 1 hr 7 mins | $1.42 |
| 21 | RFMix | 0 days 4 hrs 23 mins | $5.52 |
| 21 | Tractor | 2 days 19 hrs | $87.50 |
| 21 | Generate Final VCF | 0 days 1 hr 13 mins | $1.60 |
| 22 | Eagle | 1 day 3 hrs | $33.95 |
| 22 | Split vcf (cohort) | 0 days 3 hrs 58 mins | $5.00 |
| 22 | Split vcf (reference) | 0 days 1 hr 4 mins | $1.35 |
| 22 | RFMix | 0 days 4 hrs 44 mins | $5.95 |
| 22 | Tractor | 2 days 15 hrs | $80.75 |
| 22 | Generate Final VCF | 0 days 1 hr 19 mins | $1.71 |
